# Integrative multiomics analysis of neointima proliferation in human saphenous vein: implications for bypass graft disease

**DOI:** 10.1101/2023.11.14.567053

**Authors:** David Kim, Brandee Goo, Hong Shi, Philip Coffey, Praneet Veerapaneni, Ronnie Chouhaita, Nicole Cyriac, Ghaith Aboud, Stephen Cave, Jacob Greenway, Rohan Mundkur, Samah Ahmadieh, Ragheb Harb, Mourad Ogbi, David J. Fulton, Yuqing Huo, Wei Zhang, Xiaochun Long, Avirup Guha, Ha Won Kim, Yang Shi, Robert D. Rice, Dominic R. Gallo, Vijay Patel, Richard Lee, Neal L. Weintraub

## Abstract

**Introduction:** Human saphenous veins (SV) are widely used as grafts in coronary artery bypass (CABG) surgery but often fail due to neointima proliferation (NP). NP involves complex interplay between vascular smooth muscle cells (VSMC) and fibroblasts. Little is known, however, regarding the transcriptomic and proteomic dynamics of NP. Here, we performed multi-omics analysis in an *ex vivo* tissue culture model of NP in human SV procured for CABG surgery.

**Methods and results:** Histological examination demonstrated significant elastin degradation and NP (indicated by increased neointima area and neointima/media ratio) in SV subjected to tissue culture. Analysis of data from 73 patients suggest that the process of SV adaptation and NP may differ according to sex and body mass index. RNA sequencing confirmed upregulation of pro-inflammatory and proliferation-related genes during NP and identified novel processes, including increased cellular stress and DNA damage responses, which may reflect tissue trauma associated with SV harvesting. Proteomic analysis identified upregulated extracellular matrix-related and coagulation/thrombosis proteins and downregulated metabolic proteins. Spatial transcriptomics detected transdifferentiating VSMC in the intima on the day of harvesting and highlighted dynamic alterations in fibroblast and VSMC phenotype and behavior during NP. Specifically, we identified new cell subpopulations contributing to NP, including *SPP1^+^, LGALS3^+^* VSMC and *MMP2^+^, MMP14^+^* fibroblasts.

**Conclusion:** Dynamic alterations of gene and protein expression occur during NP in human SV. Identification of the human-specific molecular and cellular mechanisms may provide novel insight into SV bypass graft disease.

## Introduction

Human saphenous veins (SV) are the most commonly employed conduits in CABG surgery (estimated 400,000 each year in the US)^1^. Despite their widespread use, over 50% of all SV grafts fail within 10 years of implantation^2–4^, predominantly due to progressive neointima proliferation. Clinical studies have demonstrated a positive association between SV graft failure and mortality in patients undergoing CABG^5^. However, treatment of SV graft disease is clinically challenging and often requires complex interventional procedures or re-do CABG surgery, which is associated with high morbidity and mortality. To date, no medical therapies have been identified to effectively mitigate the process of neointima proliferation in SV. Prior research has identified a number of novel biochemical and molecular pathways that may contribute to SV graft failure, but thus far, the findings from these studies have not been successfully translated into clinical practice. This illustrates the critical need to better understand the mechanisms of neointima proliferation in SV in order to identify strategies to prevent SV graft failure.

The process of surgically harvesting the SV and preparing them for surgery imposes tissue trauma that is thought to serve as a trigger for neointima proliferation^6,7^. Vascular smooth muscle cells (VSMC), the major contractile cells of the vascular wall, undergo phenotype switching in response to vascular injury, leading to dedifferentiation, migration, aberrant proliferation and loss of contractile function^8^. Adventitial fibroblasts likewise have been reported to convert to myofibroblasts and migrate toward the subendothelial space following vascular injury, contributing to neointima proliferation^9^. Fibroblasts and VSMC isolated from SV have been studied *in vitro* and may provide insight into mechanisms and clinical factors associated with SV graft failure. In this regard, Kenagy et al reported that proliferative responses of VSMC and fibroblasts isolated from SV correlated with SV graft stenosis in patients undergoing bypass surgery for peripheral artery disease^10^, suggesting that the propensity to develop SV graft disease may in part be related to intrinsic properties of the VSMC and fibroblasts contained therein.

Large clinical studies have demonstrated sex disparities wherein fewer CABG surgeries are performed in women, who also experience higher perioperative mortality and worse long-term outcomes following CABG surgery, than men^11^. Additionally, an “obesity paradox” has been reported wherein overweight to moderately obese patients exhibit reduced perioperative mortality and better intermediate-term survival rates as compared to their lean counterparts^12^. The role of SV graft failure in mediating these disparities is unknown, and the influence of sex and body mass index (BMI, an indicator of obesity) on neointima proliferation in human SV has not been systematically investigated.

Recently, multiomic approaches have been employed to investigate the molecular mechanisms and cellular dynamics of neointima proliferation^13^; however, such studies cannot be readily conducted in humans, given the limited ability to procure human blood vessels. Here, we took advantage of an established tissue culture model of neointima proliferation in human SV, wherein segments of SV harvested during CABG surgery are placed in tissue culture containing growth medium, which induces neointimal proliferation over a ∼7 day period^14^. Utilizing multi-omics analyses, we detected dynamic molecular and cellular changes associated with neointima proliferation in human SV. Furthermore, we identified potential novel cell subpopulations, *MMP2^+^/MMP14^+^*fibroblasts and *LGALS3^+^/SPP1^+^* VSMC, which were not previously linked to neointima proliferation. In addition, our data suggest that mechanisms of SV adaptation and neointima remodeling may differ according to sex and BMI. Our findings provide novel insights into mechanisms of neointimal proliferation in human blood vessels and may be pertinent to SV graft disease pathogenesis and the clinical factors influencing outcomes after CABG surgery.

## Methods

### Human SV *ex vivo* culture

Collection and culturing of human SV method was adapted from our previous protocol^14^. In brief, freshly collected SV samples were obtained from patients undergoing CABG and immediately transported to the laboratory. The SV segments were placed in sterile PBS buffer and transversely cut into 3 mm segments, some of which were stored at -80° C for subsequent analyses (Day 0). Remaining segments were cultured for 7 days (Day 7) in RPMI 1640 containing NaHCO_3_ 2 g/L, penicillin/streptomycin (100 IU/100 μg per ml), L-glutamine 4 ml/L, and 30% fetal bovine serum at 37°C in 5% CO_2_. Media was replaced every other day, and then, segments were stored at -80° C for subsequent analyses. Collection of the human SV and related experiments were designated as exempted from human subjects research by the Institutional Review Board (IRB) at Augusta University.

### Bulk RNA sequencing in SV and data analysis

Total RNA from six biological replicates of Day 0 and Day 7 human SV with RNA integrity number (RIN) greater than 8 was sent to Genewiz for preparation into RNA-Seq libraries. Polyadenylated RNA was sequenced at a depth of 20 million reads per replicate using Illumina HiSeq 2500 system, using a 150 bp paired-end protocol (Illumina). Raw reads were trimmed using cutadapter version 4.1, and the processed reads were mapped to the GRCh38/hg38 genome assembly by using STAR version 2.7.0^15^.

DESeq2 version 1.22 was used to generate normalized read counts and conduct differential gene expression analysis with a false discovery rate (FDR) threshold of less than 0.05 using R studio version 494 (https://posit.co/download/rstudio-desktop/) and public server of usegalaxy.org^16,17^. PCA and volcano plot was created through ggplot2. Gene ontology analysis and gene-network graph were performed and created through clusterProfiler version 3.0.4^18^.

### Liquid chromatography tandem mass spectrometry (LC-MS/MS) analysis

Protein lysates from three biological replicates of Day 0 and Day 7 SV were submitted to the Proteomics Core at Augusta University. Protein digestion and mass spectrometry were performed as previously described^19^. Briefly, protein lysates were separated through Ultimate 3000 nano-UPLC system (Thermo Scientific) and run on an Orbitrap Fusion Tribrid mass spectrometer (Thermo Scientific). Raw data were analyzed using Proteome Discoverer (v1.4, Thermo Scientific) and searched against the UniProt human protein database. Peptide-spectrum match (PSM) values, a total number of identified peptide spectra matched for the protein, were log-transformed to normalize the dataset, and missing values were imputed through Amica version 3.0.1^20^. After normalization of the dataset, differential protein expression analysis was performed with threshold set at FDR of less than 0.05. Gene ontology analysis and gene-network graph were performed and created through clusterProfiler version 3.0.4^18^. For labeled proteomics, proteins of interest were labeled using stable isotopes, and the same protein lysates were subjected to liquid chromatography coupled with mass spectrometry as previously described. The labeled peptides were quantified to identify differential protein expression across experimental conditions.

### Visium Spatial Transcriptomics

FFPE Visium Spatial Transcriptomics data was generated according to the manufacturer’s protocol. Briefly, FFPE sections from three biological replicates of Day 0 and Day 7 SV with DV200 greater than 50% were mounted on a glass slide and sent to the Advanced Genomics Core at the University of Michigan. After H&E images were taken, the resulting cDNA libraries were sequenced on a NovaSeq 6000 system (Illumina). Demultiplexed FASTQ files were converted to count matrices using SpaceRanger version 1.3.1 (10X Genomics), and the resulting datasets were mapped to GRCh38. The mapped sequencing data was processed with the Visium Human Transcriptome Probe Set v 2.0 (10X Genomics), and spot annotations were performed in Loupe Browser 5 (10X Genomics). Seurat Object for the sequenced samples was constructed from the count matrices and spot annotations. For downstream analysis, spots less than 500 unique molecular identifiers (UMIs) and greater than 5% mitochondrial reads were considered spots of low quality reads, and subsequently excluded from the dataset. Subsequently, high-quality spots were normalized by SCTransform in the Seurat package^21,22^. All datasets were merged, and dimensionality reduction was conducted on the pooled dataset using the Seurat package, and around 5000 spots were clustered. The cellular identities of each cluster were predicted using previously published single-cell transcriptomic signatures, in addition to the tissue morphology underneath the spots for the human SV^23,24^. Genes with FDR of less than 0.05 were differentially expressed using Benjamini-Hochberg method. Top differentially expressed genes (DEGs) were selected based on the absolute value during the DE analysis. Gene ontology analysis was performed as previously described, and pseudotime trajectory analysis was performed using Monocle3^25^.

### Quantitative PCR

Total RNA was extracted from human SV with QIAzol Lysis Reagent, and purified with miRNeasy Mini Kit (Qiagen). Real time quantification of mRNA levels of the genes of interest was performed using Brilliant II SYBR Green QPCR Master Mix (Agilent Technologies) per manufacturer’s instructions. Normalized Ct values were subjected to statistical analysis and fold change was calculated by ΔΔ Ct method as described previously, normalized against 18S^26,27^. Primer sequences are listed in Table S1.

### Western Blotting

Protein extraction and Western blotting were performed as described previously^26^. Antibodies used are listed in Table S2.

### Immunohistochemistry

Paraffin-embedded SV tissue sections were stained with hematoxylin and eosin (H&E), Verhoeff van Gieson (VVG), α-SMA, VE-cadherin, and DAB Substrate (Vector Labs) kits were used for visualization as previously described^26^. Quantification of intima, media, adventitia and total vessel area of SV was analyzed with ImageJ software (https://imagej.net) using freehand selection tool on VVG stained slides. The number of elastin breaks was counted in three sections per tissue sample to quantify elastin degradation. To quantify the immunostaining data, stained area in the intimal and medial region was analyzed using ImageJ software. Antibodies used are listed in Table 2.

**Table 1.**
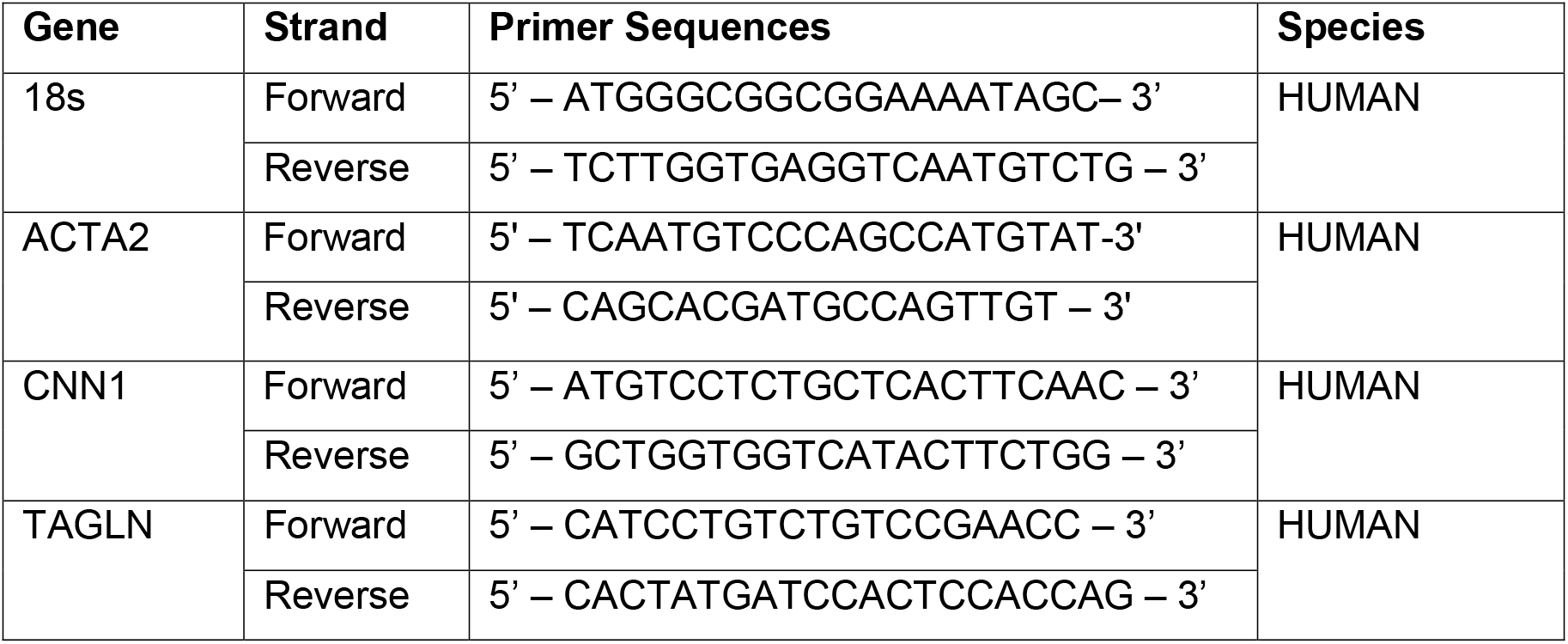
Sequences for qPCR primers.

**Table 2.**
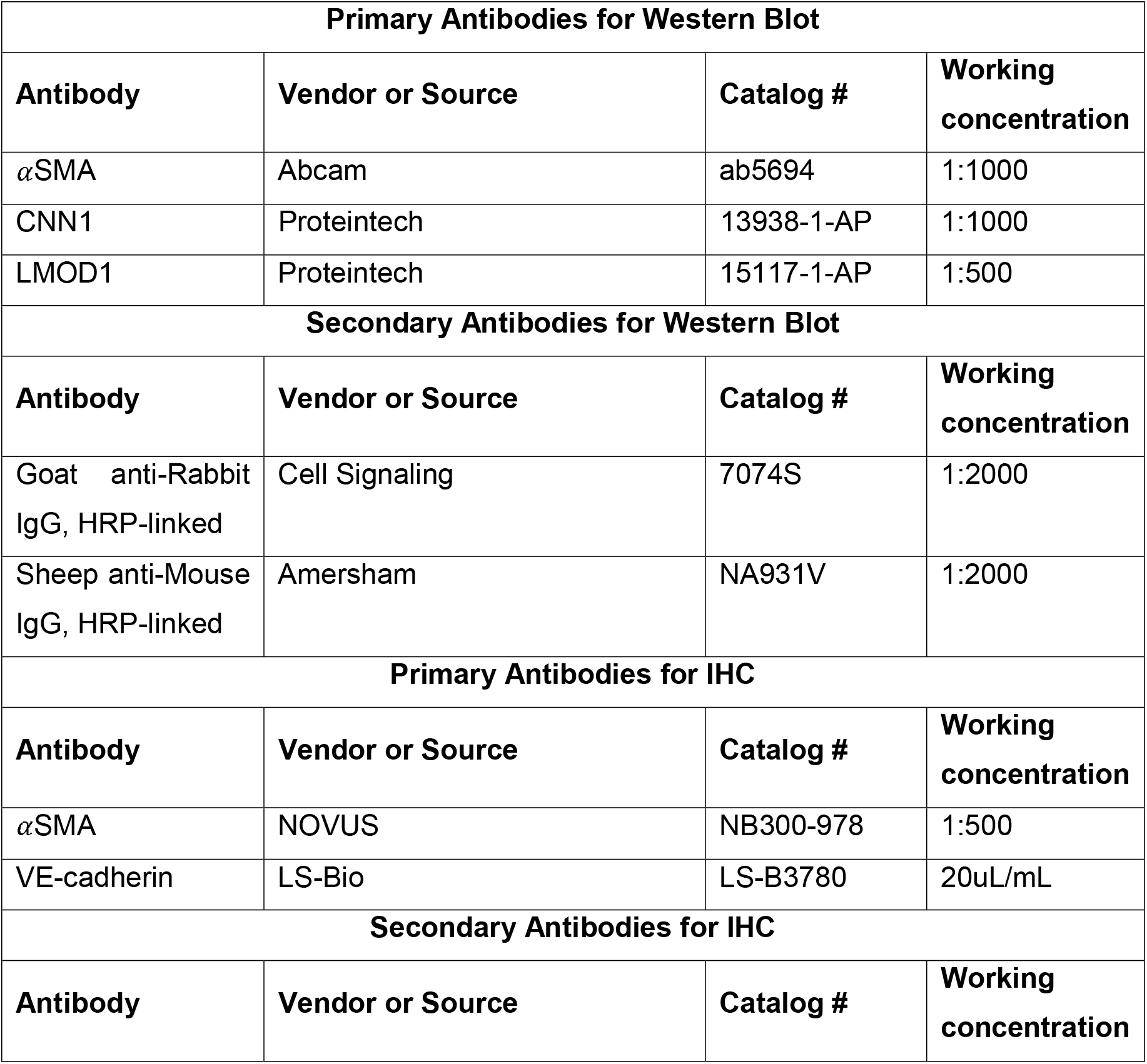
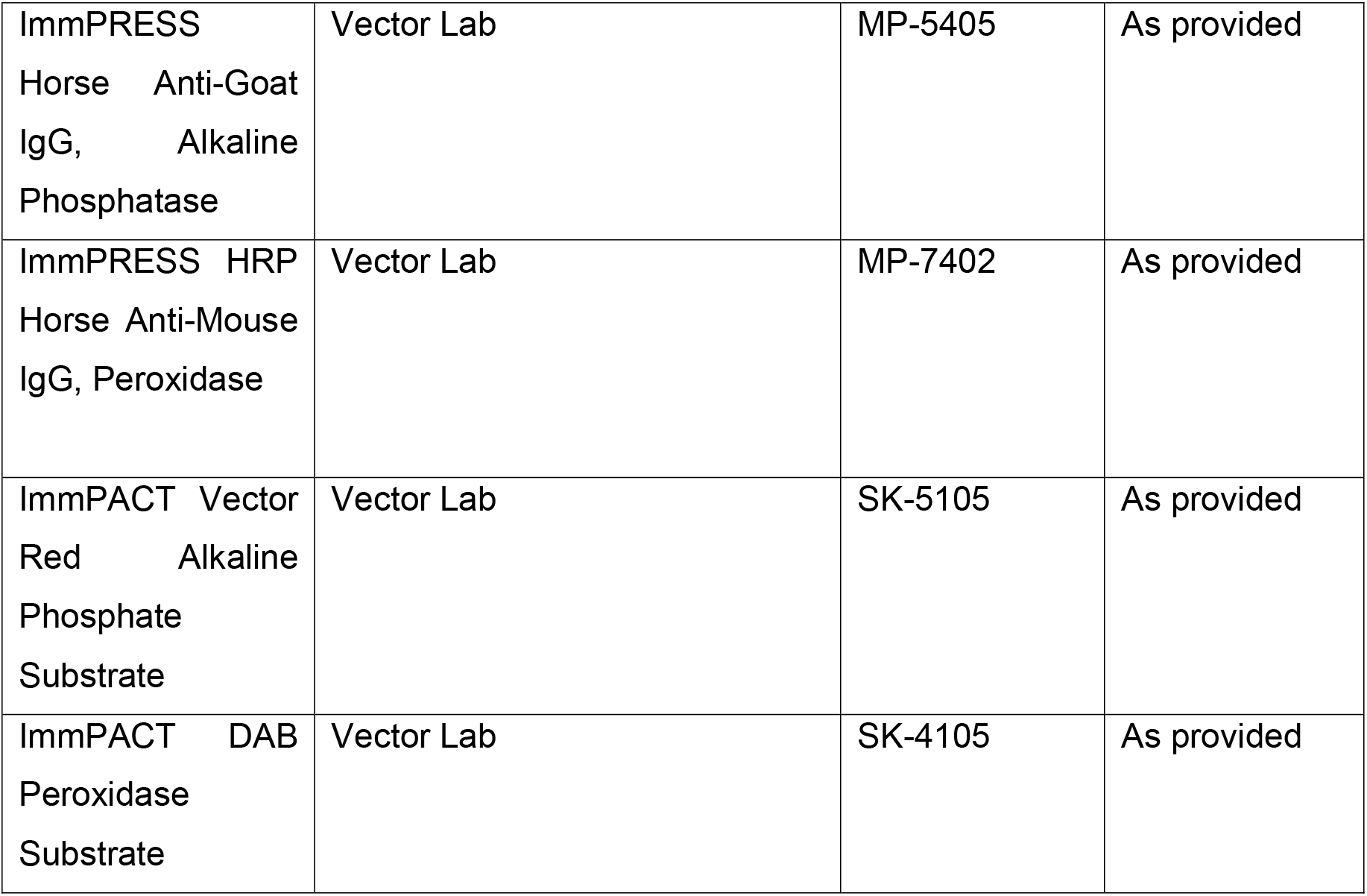
Antibodies for Western Blot and Immunohistochemistry.

## Results

### Human SV incubation model

Human SV segments were obtained in patients undergoing CABG surgery as previously reported^14^. The experimental workflow is depicted in **Figure 1A**. Veins were either processed immediately for investigation (Day 0) or maintained in a cell culture incubator *ex vivo* for 7 days (Day 7), after which they were processed for investigation using identical procedures. Histological examination (H & E staining) confirmed significant neointima proliferation in SV at Day 7 compared to Day 0 (**Figure 1B, left panel**).

**Figure 1.**
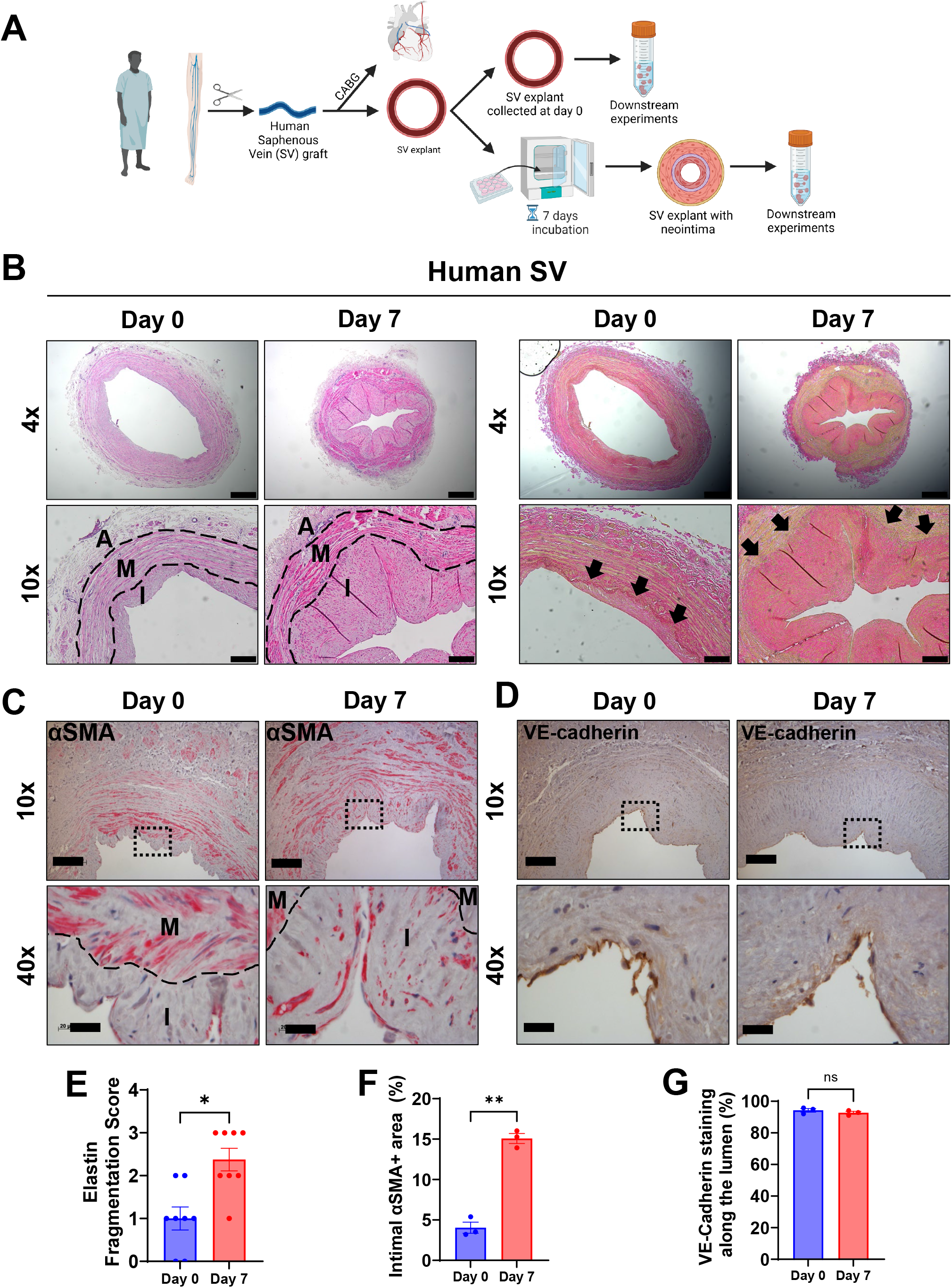
Human saphenous vein (SV) explants undergo neointima formation ex vivo. A, Schematic of ex vivo model of neointima formation in human SV. B, Representative image of H&E (left) and VVG (right) staining in unincubated (Day 0) and incubated (Day 7) human SV. Scale bar: 500 μm (4x) and 200 μm (10x). C, Representative images of α-Smooth Muscle Actin (αSMA; red) staining in Day 0 and Day 7 SV. Scale bar: 200 μm (10x) and 20 μm (40x). D, Representative images of VE-cadherin (brown) staining in Day 0 and Day 7 SV. Scale bar: 200 μm (10x) and 20 μm (40x).E-G, Quantification of elastin fragmentation score (E, from VVG staining, n=8), αSMA positive area in the intima (F, n=3), and VE-cadherin staining along the luminal lining (G, n=3). Data are mean±SEM by paired t-test (n=3).I, intima; M, media, A adventitia.

### Key aspects of neointima proliferation in the *ex vivo* incubation model

The *ex vivo* model of neointima proliferation in human SV was further characterized histologically. We first performed VVG staining to detect elastin fibers in human SV. VVG staining demonstrated that the internal elastic lamina was fully intact at Day 0, whereas significant disruption and fragmentation was apparent at Day 7 (**Figure 1B, right panel,** quantified in **1E**).

To assess VSMC accumulation in the intima, a key aspect of neointima proliferation, we performed α-SMA staining on Day 0 and Day 7 SV. As expected, SV at Day 0 demonstrated little α-SMA positive staining within the intima, while α-SMA staining was readily apparent in the proliferated neointima at Day 7 (**Figure 1C, 1F**). Additionally, VE-cadherin immunostaining demonstrated the consistent presence of endothelial cells (EC) along the luminal surface in both Day 0 and Day 7 SV, suggesting preserved EC integrity in this model (**Figure 1D**, quantified in **1G**).

### Neointima quantification and separation by sex

Next, we extended the *ex vivo* incubation model of neointima proliferation to include 73 paired SV samples from individual patients. Quantification of the neointima area and neointima-to-media ratio demonstrate statistically significant increases at Day 7 compared to Day 0 (**Figure 2A**). Individual (paired) results from the same patients are shown in **Figure 2B**; while there was variability in the magnitude of the response, the majority of SV from individual patients exhibited neointima proliferation at Day 7.

**Figure 2.**
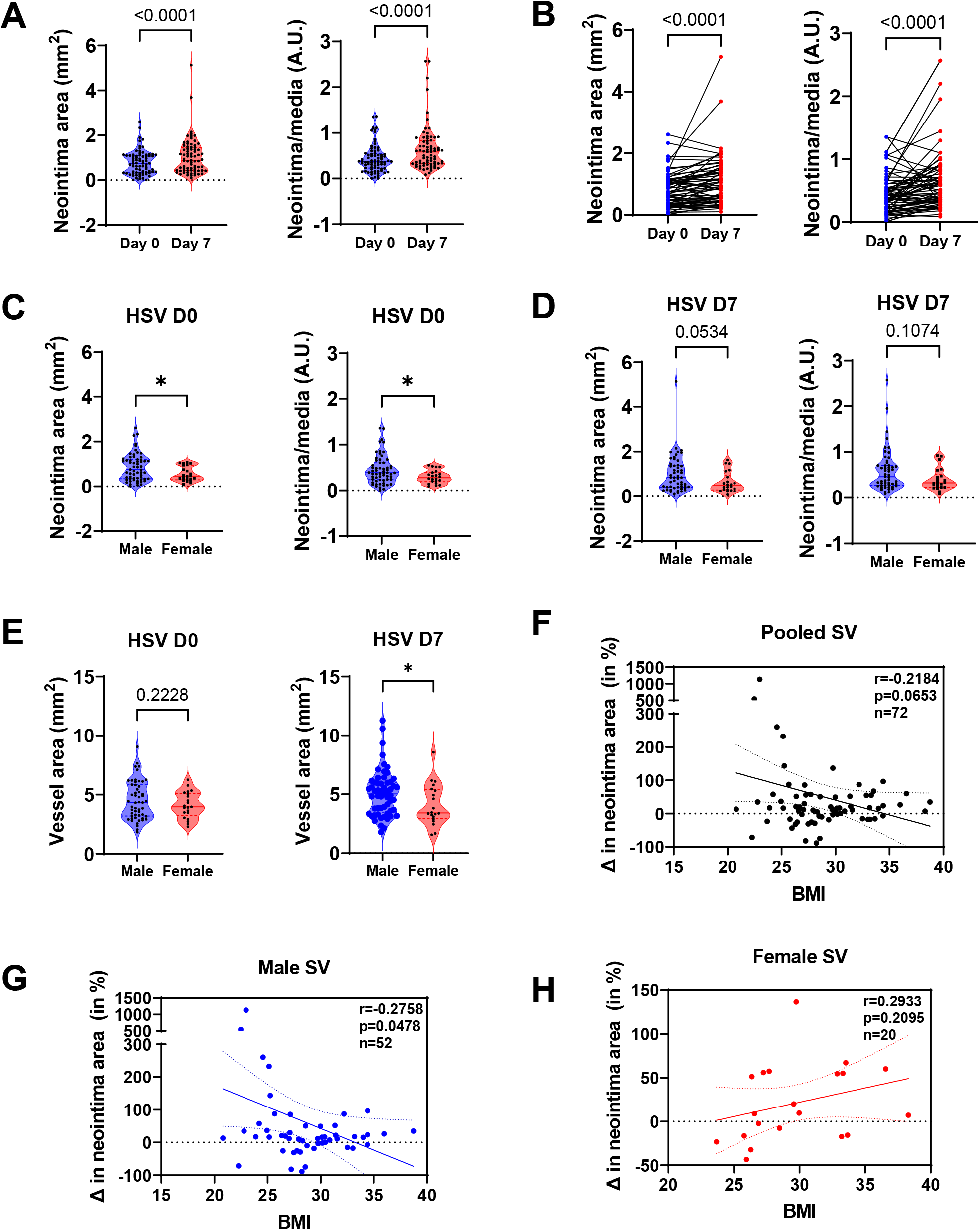
Sex differences in neointima induction in relation to BMI in human SV **A,** Quantification of neointima area and neointima-to-media ratio of Day 0 and Day 7 human SV (paired). Data are mean±SEM by Wilcoxon signed-rank test (n=73). **B,** Before-after line graphs of neointima area and neointima-to-media ratio in SV at Day 0 and Day 7 (paired). Data are mean±SEM by Wilcoxon signed-rank test (n=73). **C,D** Quantification of neointima area and neointima-to-media ratio of Day 0 (C) and Day 7 (D) human SV, separated by sex. Data are mean±SEM by Mann-Whitney test (Male: n=53; female: n=20). **E,** Quantification of total vessel area of Day 0 (**left**) and Day 7 (**right**) human SV, separated by sex. Data are mean±SEM by Mann-Whitney test (Male: n=53; female: n=20). **F-H,** Scatterplot of BMI in relation to the change of neointima area between Day 0 and Day 7 SV in the entire cohort (n=72) and in males (**G**, n- 52) and females **(H**, n=20).

SV tend to be smaller in diameter in females than in males^28^. Moreover, SV graft disease in females was reported to differ pathologically from that in males, exhibiting a more prominent cellular fibrous tissue component^29^. Thus, we analyzed the SV data separately by sex. As expected, far more specimens were collected from males (n=53) than females (n=20). Interestingly, at Day 0, SV from females had significantly less neointima compared to males, but this difference was diminished by Day 7. We also compared total vessel area from female versus male patients at Day 0 and Day 7. While no significant differences were noted at Day 0, total SV area was significantly smaller in women compared to men at Day 7.

Overweight and mildly obese individuals have been reported to exhibit improved intermediate term survival after CABG^30^. Therefore, we separated the SV data by BMI. In both Day 0 and 7 SV, we did not detect an association between neointima area and BMI. We then quantified the change in neointima between Day 0 and Day 7 (an indicator of neointima proliferation), which showed a strong trend towards a negative association in the entire cohort (**Figure 2F**). Notably, when we separated the BMI data by sex, SV from males exhibited a significant negative association between change in neointima and BMI, while SV from females trended towards a positive association, although the numbers were small and statistically non-significant. These findings suggest that mechanisms of SV adaptation and neointima remodeling may differ according to sex and BMI.

### RNA sequencing and multiomics profiling of saphenous vein

Workflow for multiomics experiment is depicted in **Figure 3A**. To examine the global transcriptomic changes occurring during neointima proliferation, bulk RNA-sequencing was first performed in patient-matched Day 0 and Day 7 SV. Principal component analysis (PCA) plot depicted divergent and independent clustering of Day 0 and Day 7 SV, indicating distinct gene expression profiles between the two time points (**Figure 3B**).

**Figure 3.**
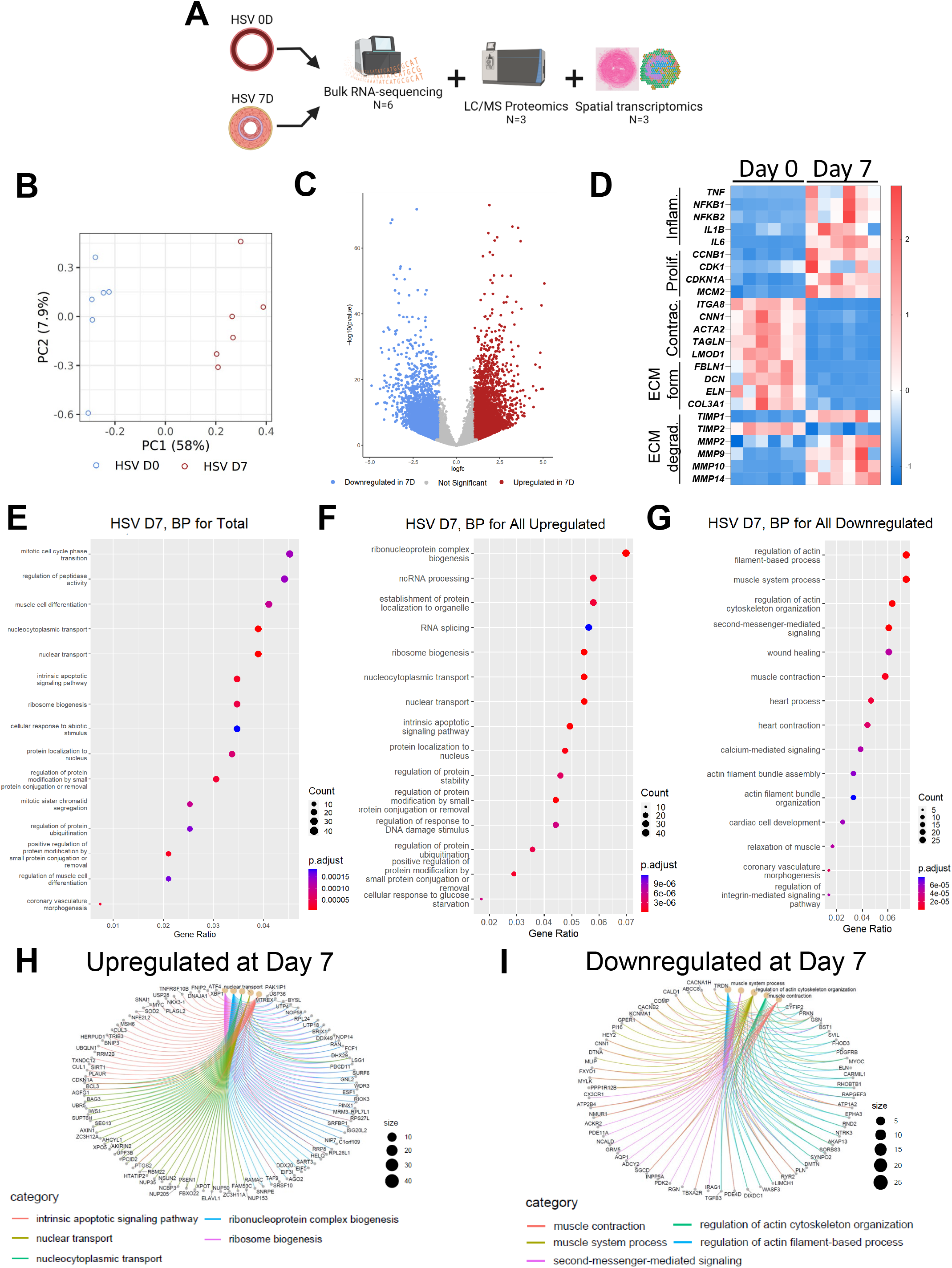
Multi-omics profiling of neointima formation in human SV **A,** Schematic of workflow for the multi-omic study. **B,** PCA plot from bulk RNA-sequencing depicting Day 0 (**blue**) and Day 7 (**red**) SV. **C,** Volcano plot comparing the differentially expressed genes between Day 0 and Day 7 SV. Each dot represents significantly increased (red) or decreased (blue) genes in Day 7 compared to Day 0 SV. **D,** Heatmap of selected genes which were highly differentially expressed between Day 0 and Day 7 SV. **E-G,** Top GO terms of all (**E**), upregulated (**F**) and downregulated (G) DE genes (Fold Change <0.5 OR >2, FDR < 0.01). **H**, Gene-plot network reconstruction of top GO terms from panel F. **I,** Gene-plot network reconstruction of top GO terms from panel G.

A total of 36969 genes were detected, with 2416 upregulated genes and 1967 downregulated genes in Day 7 SV compared to Day 0 SV (Fold Change <0.5 OR >2, FDR < 0.01, **Figure 3C**). As expected, pro-inflammatory (*TNF, NFKB1, NFKB2, IL1B, IL6*) and proliferation-related (*CCNB1, CDK1, CDKN1A, MCM2*) genes were significantly increased in Day 7 compared to Day 0 SV. Additionally, smooth muscle differentiation-related genes (*ITGA8, CNN1, ACTA2, TAGLN, LMOD1*) and extracellular matrix forming genes (*FBLN1, DCN, ELN, COL3A1*) were significantly downregulated in Day 7 compared to Day 0 SV (**Figure 3D**). To interrogate the lesser known molecular and cellular mechanisms associated with neointima formation, pathways and gene ontology (GO) enrichment analyses were performed. At the global level, GO analysis revealed upregulation of genes which govern RNA-related processes, intrinsic apoptotic signaling pathway and protein ubiquitination in Day 7 SV (**Figure 3E, 3F**). By contrast, genes which control actin modulation, cytoskeletal organization and second messenger signaling were consistently downregulated in Day 7 SV (**Figure 3E, 3G**). Notably, no strong correlation in genes regulating extracellular matrix degradation was detected; while some matrix metalloprotease-related genes were significantly increased (*TIMP1, MMP10*), others were not significantly changed between Day 0 and Day 7 SV (**Figure 3D**).

To identify the most differentially regulated GO processes, we constructed a gene-plot network, which provides a visual representation of the interactions and relationships among genes, aiding in the comprehensive analysis of complex biological systems. This analysis suggested increased nuclear transport and ribosome biogenesis in Day 7 SV (**Figure 3H**). Taken together with the more complex reads from Day 7 SV, these data suggest that higher levels of both transcription and translation are occurring at this time during neointima formation. As expected, the gene-plot network revealed downregulation of actin-cytoskeleton modulating genes, muscle contraction genes, and second messenger signaling, similar to the GO pathway analysis at D7 (**Figure 3I**).

### Proteomic profiling of neointima proliferation

Next, we performed unlabeled LC/MS proteomics on SV lysates. A total of 3380 proteins were detected, with 55 upregulated proteins and 74 downregulated proteins in Day 7 SV compared to Day 0 SV (Fold Change <0.5 OR >2, p-value < 0.05). Similar to the RNA-seq data, PCA plot of the proteomics samples demonstrated distinct clustering of proteins in Day 0 versus Day 7 SV, although samples within the same group showed a wide degree variability (**Figure 4A**), as was likewise observed with respect to neointima formation (**Figure 2B**). Interestingly, the top upregulated proteins in Day 7 SV represented a mixture of both ECM-producing and ECM-degrading proteins (COL4A1, MMP2, FBN1, EMILIN1), while the top downregulated proteins represented proteins which regulate metabolic processes, including mitochondrial metabolism (IDH2, ACO2, HADHA, ALDH2) (**Figure 4B**).

**Figure 4.**
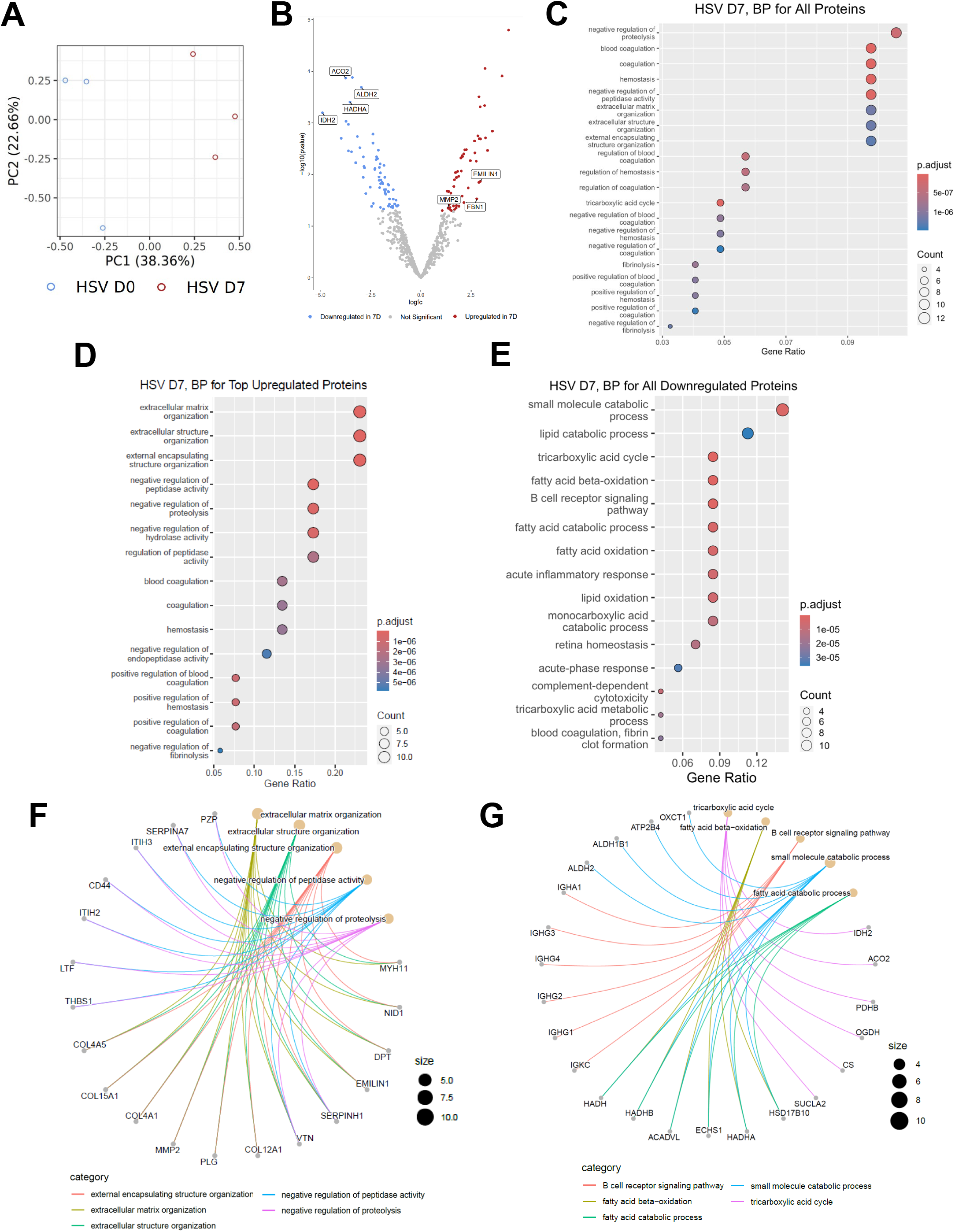
Proteomic profiling of neointima formation **A,** PCA plot from untargeted LC/MS proteomics depicting the Day 0 (**blue**) and Day 7 (**red**) SV. **B,** Volcano plot comparing the differentially expressed proteins between Day 0 and Day 7 SV. **C-E,** Top GO terms of all (**C**), upregulated (**D**) and downregulated (**E**) DE proteins (Fold Change <0.5 OR >2, p-value < 0.05). **F**, Gene-plot network reconstruction of top GO terms from panel D. **G,** Gene-plot network reconstruction of top GO terms from panel E.

GO and gene-plot network analyses showed extracellular matrix organization and negative regulation of proteolysis/peptidase activity to be amongst the most significant terms, consistent with increased ECM-producing proteins in Day 7 SV (**Figure 4C, 4D, 4F**). Interestingly, other GO terms included blood coagulation and hemostasis, suggesting that changes in protein expression occurring during the early phase of SV remodeling may promote thrombosis (**Figure 4D**). Top downregulated protein GO analysis terms included small molecule catabolic process, tricarboxylic acid cycle, and fatty acid beta-oxidation, indicating significant metabolic dysfunction during neointima formation (**Figure 4C, 4E, 4G**). Additional top downregulated terms included decrease of B cell receptor signaling pathway, in particular IgG heavy chains (IGHA1, IGHG1, IGHG2, IGHG3, IGHG4) (**Figure 4G**). Using this unlabeled proteomics approach, expression of most VSMC contractile proteins was not significantly reduced, and one contractile protein, MYH11, appeared to exhibit an increase in expression in Day 7 SV.

### Spatial transcriptomic analysis suggests dynamic alterations in fibroblast and VSMC phenotype and behavior

Given the dynamic transcriptional and translational changes in SV during neointima formation, we next performed spatial transcriptomic analysis to interrogate gene expression locally within blood vessel wall at Day 0 and Day 7. Representative images from this experiment are shown in **Figure 5A**. Merged UMAP plot of the barcoded spots demonstrate a total of 8 clusters, with large separation between Day 0 and Day 7 SV, and minimal overlap of spots (**Figure 5B**). Day 0 SV contains 5 separate clusters with distinct spatial localization and gene expression profiles. Transdifferentiating VSMC expressing both VSMC-specific and proliferation markers (i.e., *MYH11, CNN1, MKI67*, and *PCNA*) were detected in the intima (**Figure 5A**). Contractile VSMC with the highest expression of VSMC markers were detected in the medial layer. Unexpectedly, however, these contractile VSMC represent a minority of cells in the medial layer, with fibroblasts being the dominant cell type in both the media and adventitia at Day 0. Day 7 SV exhibited almost no contractile VSMC, and instead showed several novel clusters which were not detected in Day 0 SV, including synthetic and osteogenic-like VSMC. Interestingly, secretory fibroblasts, with high expression of *FBLN1, FBN,* and *DCN*, were dominant in Day 0 SV and replaced by degradative fibroblasts with high expression of metalloproteases, including *MMP2, TIMP1,* and *MMP14* in Day 7 SV (**Figure 5B, 5C**). We did not detect clusters representing EC in either Day 0 or Day 7 SV.

**Figure 5.**
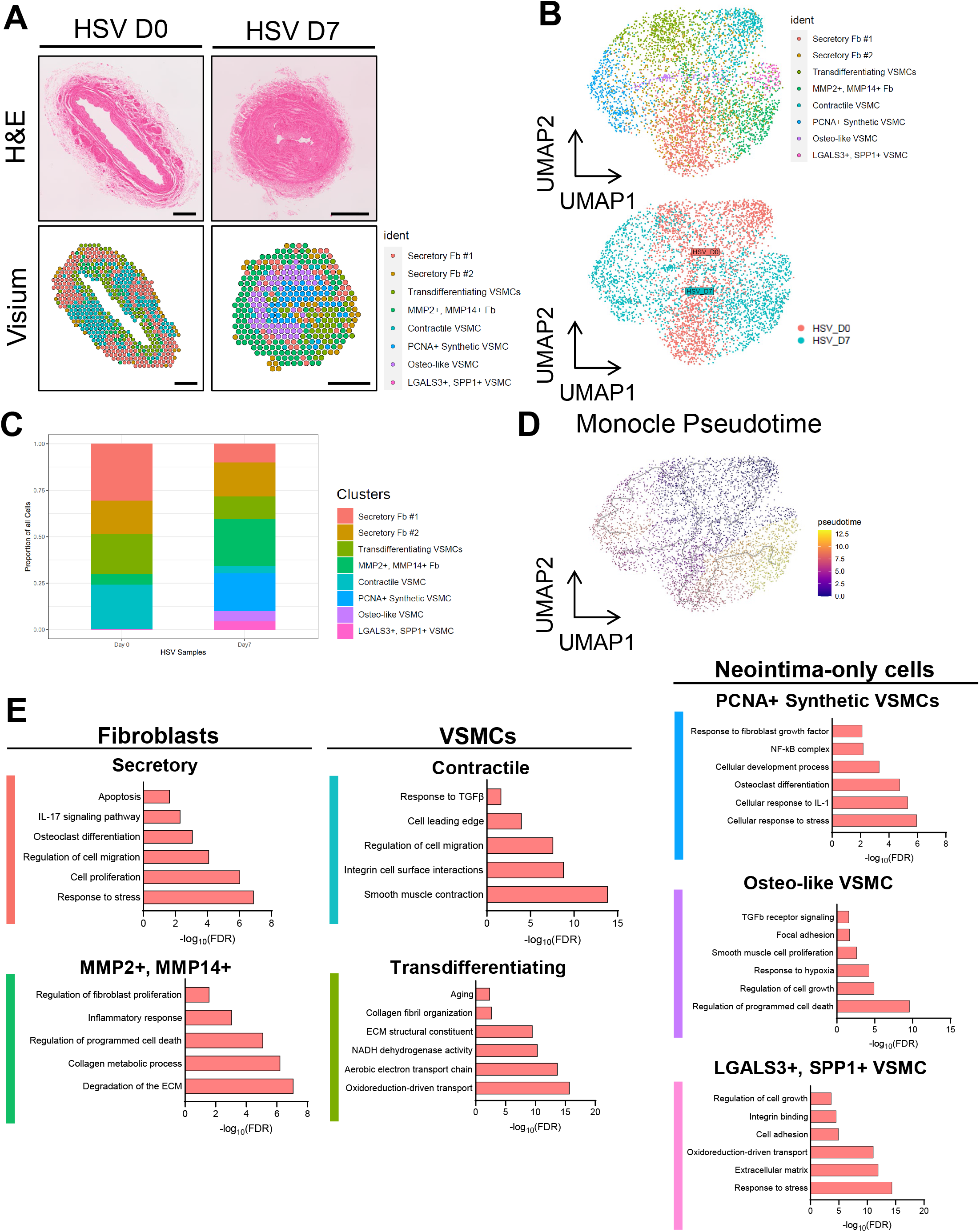
Spatial transcriptomics analysis of human SV during neointima proliferation **A,** Representative image of H&E (**top**) and 10X Visium spatial transcriptomics (**bottom**) in Day 0 and Day 7 SV. Scale bar: 500 μm. **B,** Uniform Manifold Approximation and Projection (UMAP) plot of barcoded spots identified by Seurat (**top**), and UMAP plot split by condition (Day 0 vs Day 7 SV, **bottom**). **C**, Percentage change of each cluster during neointima proliferation and the contribution of each cluster to Day 0 and Day 7 SV. **D**, Trajectory analysis of the merged UMAP plot for Day 0 and Day 7 SV. **E**, Gene Ontology (GO) enrichment for differentially expressed genes within each cluster.

Pseudotime trajectory analysis of the merged UMAP confirms the presence of contractile and transdifferentiating VSMC in Day 0 SV, with progression into synthetic VSMC, secretory fibroblasts and eventually *MMP2+, MMP14+* fibroblasts (**Figure 5D**). GO analysis showed distinct processes which align with the cluster descriptions. Notably, cellular responses to stress and apoptosis were the highest represented pathways in intimal VSMC clusters, which align with the findings from bulk RNA-sequencing (**Figure 5E)**. GO terms of transdifferentiating and *LGALS3^+^, SPP1^+^* VSMC clusters included oxidoreduction-driven transport and extracellular matrix, which were also GO terms enriched in both the bulk RNA-seq and proteomics data. Furthermore, GO terms of the secretory fibroblasts highly present in Day 0 SV included proliferation and response to stress, while GO terms of *MMP2^+^, MMP14^+^* fibroblasts present in Day 7 SV included degradation of ECM and inflammatory response. Taken together, these data demonstrate that the VSMC and fibroblast clusters undergo dynamic fate changes during neointima proliferation in human SV.

### Global VSMC transcriptional changes during neointima formation in human SV

To assess global VSMC transcriptional changes during neointima formation in SV, we first examined VSMC contractile genes through spatial transcriptomics and present the results in violin plots (**Figure 6A, 6B**). These findings confirm reduced VSMC contractile gene expression at the whole tissue level. Our unlabeled proteomics analysis did not show significant reductions in VSMC proteins (**Figure 4B**). Thus, we performed targeted analysis of VSMC contractile proteins, including calponin and transgelin, which showed that the area under the curve is significantly less in Day 7 SV than in Day 0 SV, while beta-actin showed no significant difference (**Figure 6D**). Western blot of these VSMC proteins showed significant decrease in contractile proteins at day 7, as indicated in **Figure 6E, 7F**^31^. Interestingly, spatial transcriptomics data also showed that ECM-forming genes (e.g., FBLN1, DCN, COL3A1 and ELN) were downregulated whereas ECM-degrading genes (e.g., MMP2, MMP14) were upregulated during neointima proliferation, suggesting transcriptional changes in gene expression and phenotype switching in fibroblasts potentially linking to neointima proliferation.

**Figure 6.**
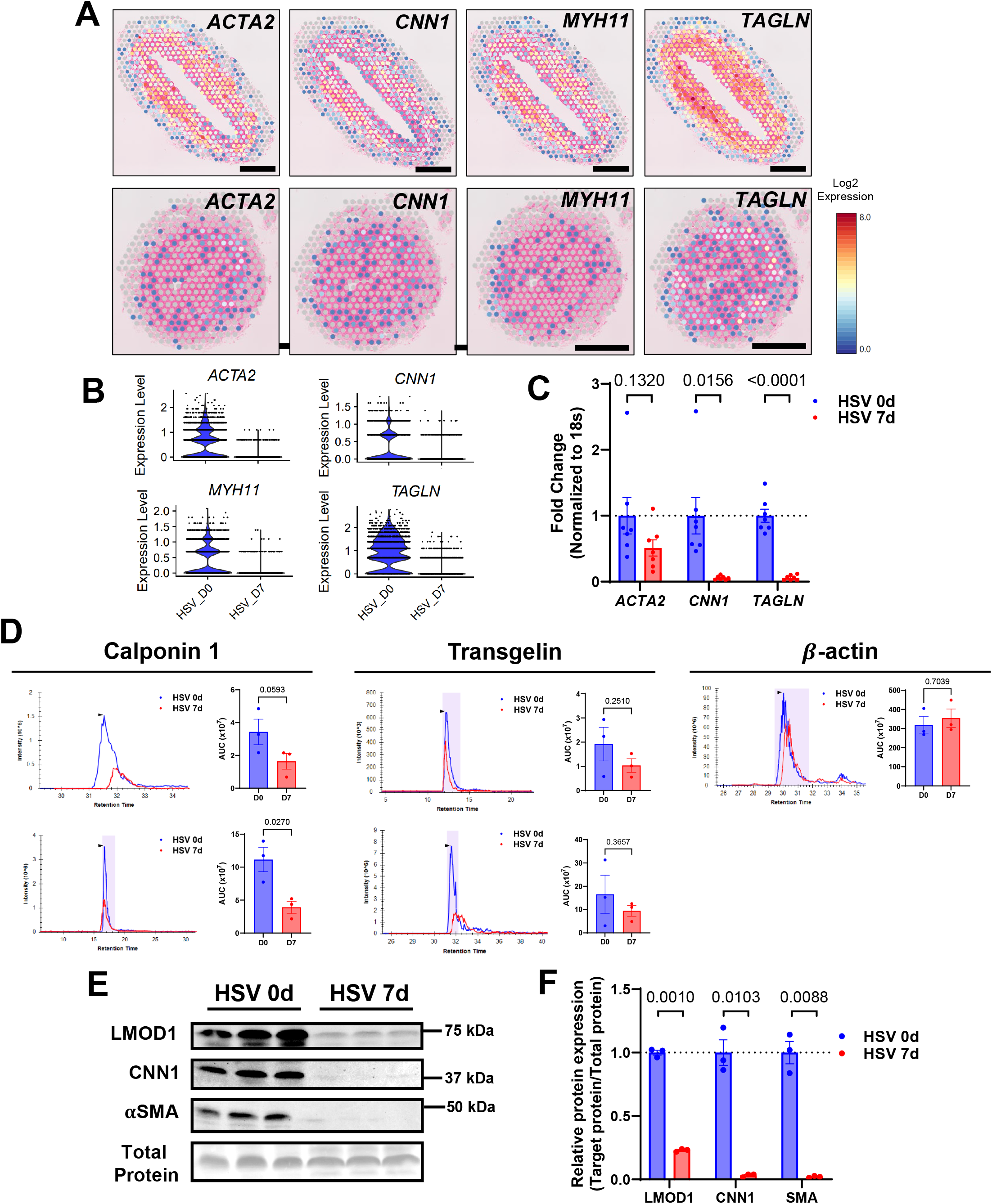
VSMC in human SV undergo de-differentiation during neointima proliferation **A,** Visium spatial transcriptomics feature plot of VSMC contractile markers (*ACTA2, CNN1, MYH11, TAGLN*) in Day 0 (**upper**) and Day 7 (**lower**) SV. Scale bar: 500 μm. **B,** Violin plot of VSMC contractile genes (*ACTA2*, *CNN1, MYH11, TAGLN*) in Day 0 and Day 7 SV (n=3). **C,** qRT-PCR of representative VSMC contractile genes (*ACTA2, CNN1, TAGLN*) in RNA from Day 0 and Day 7 SV. Data are mean±SEM by Student’s t-test (n=7). **D,** Targeted proteomics analysis of VSMC contractile proteins (Calponin 1, Transgelin) and β-actin in Day 0 and Day 7 SV. Data are mean±SEM by Student’s t-test (n=3). **E,** Western blot analysis of LMOD1, CNN1, and αSMA in Day 0 and Day 7 SV. **F,** Mean densitometric analysis for LMOD1, CNN1, and αSMA normalized to the total protein. Data are mean±SEM by Student’s t-test (n=3).

**Figure 7.**
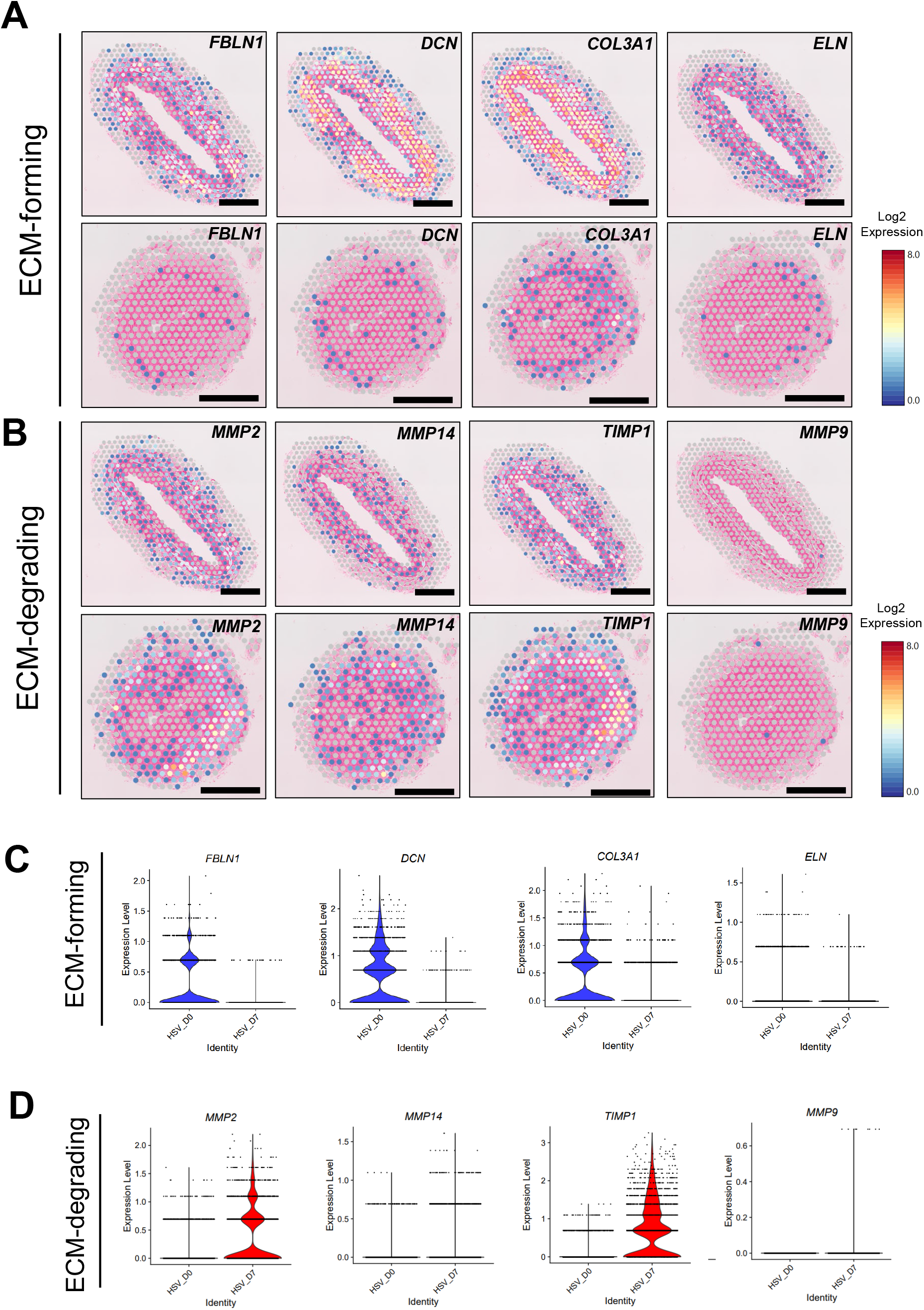
Transcriptional changes in genes regulating the extracellular matrix (ECM) in fibroblasts during neointima proliferation in human SV **A,** Visium spatial transcriptomics feature plot of ECM-forming genes (*FBLN1, DCN, COL3A1, ELN*) in Day 0 (**upper**) and Day 7 (**lower**) SV. Scale bar: 500 μm. **B,** Visium spatial transcriptomics feature plot of ECM-degradation genes (*MMP2, MMP14, TIMP1, MMP9*) in Day 0 (**upper**) and Day 7 (**lower**) SV. Scale bar: 500 μm. **C,** Violin plot of ECM-forming genes (*FBLN1, DCN, COL3A1, ELN*) in Day 0 and Day 7 SV (n=3). **D,** Violin plot of ECM-degradation genes (*MMP2, MMP14, TIMP1, MMP9*) in Day 0 and Day 7 SV (n=3).

## Discussion

Saphenous veins remain the most commonly used conduit for CABG surgery, yet 50% of vein grafts are reported to fail within 10 years of implantation^2,3^. While early vein graft failure is typically associated with technical factors and/or thrombosis, most SV fail over time due to progressive neointima proliferation^32–34^. The process of neointima proliferation in SV is thought to be initiated by injury to the vessel wall sustained during surgical harvesting, pressurization and preparation for implantation^35^. Consequently, when SV segments harvested for CABG grafting are maintained in tissue culture, they form a VSMC-rich neointima. Here, we performed a detailed characterization of neointima proliferation in human SV, and we employed multi-omics to investigate cellular dynamics and molecular mechanisms therein. These findings may provide novel insight into the pathogenesis of SV graft disease and could help to illuminate two poorly-understood disparities (sex and obesity) in patients undergoing CABG surgery.

A total of 73 paired samples from individual patients were studied on the day of harvesting (Day 0) or after 7 days (Day 7) in tissue culture to elicit neointima proliferation. In general, neointima proliferation in this model was characterized by pronounced accumulation of VSMC and degradation of elastin fibers, while the endothelial monolayer remained intact. However, we detected considerable variability in the extent of neointima present in the SV in this study, both at Day 0 and Day 7. Our findings indicate that this variability might be linked, at least in part, to sex-related distinctions, wherein SV from males exhibited a larger intima and intima/media ratio than females at Day 0. This sex difference in neointima size was less apparent at Day 7, however. Additionally, the total vessel area was similar in men and women at Day 0 but significantly smaller in women at Day 7. It should be noted that women were under-represented in this study, as has been the case with other studies involving CABG surgery, in which women typically represent 20-30% of the CABG population^36^. Nevertheless, these findings raise the possibility that early responses to injury and adaptive remodeling of implanted SV might differ in women compared with men. Women have been reported to experience a higher rate of myocardial infarction and re-vascularization after CABG surgery^37^; whether our findings might contribute to such sex-dependent differences in outcomes after CABG surgery remains to be determined.

Traditional atherosclerosis risk factors, including smoking, hypertension, diabetes and hyperlipidemia, have been associated with SV graft failure in humans^38^. However, factors which may protect against SV graft failure have not been identified. Interestingly, we observed a strong trend towards a negative association between BMI and neointima proliferation in our cohort. Breaking down the data by sex, we found that the association was significant in men, but not in women, in whom the data trended in the opposite direction (i.e., towards a positive association between BMI and neointima proliferation). Aside from the fact that this may represent another sex difference in SV remodeling, our findings may help to explain an apparent “obesity paradox” that has been reported in studies of CABG, wherein overweight to moderately obese individuals tend to exhibit better outcomes and intermediate-term survival rates compared to their leaner counterparts^12,39^. This benefit appears to be lost over time, likely due to the adverse health burden imposed by obesity and its associated risk factors (i.e., hypertension, hyperlipidemia, insulin resistance, etc.)^40^. To the best of our knowledge, very few longitudinal studies have reported bypass graft patency data, and none of them examined the relationship between BMI and graft patency, so the clinical significance of our findings remains uncertain. It should also be noted that morbid obesity is an independent risk factor for adverse outcomes following CABG; however, in this study, none of our patients exhibited a BMI of > 40 and thus were not classified as morbidly obese.

In this study, we performed a multi-omics analysis of human SV to probe the molecular and cellular mechanisms involved in neointima proliferation. We show that the process of neointima proliferation is highly dynamic, with nearly half of the entire transcriptome exhibiting changes in expression at Day 7 as compared to Day 0 (p-adjust < 0.05). Bulk RNA-seq analysis demonstrated a number of upregulated genes and pathways during neointima proliferation, many of which (i.e., inflammation, cell proliferation) have been reported in other model systems^41,42^. In addition, we detected downregulation of genes associated with VSMC differentiation, which has also been reported previously^43^. Gene ontology analysis demonstrated that during neointima proliferation, pathways associated with intrinsic apoptosis and response to DNA damage stimulus are highly upregulated. These processes occur in tandem with increased transcription and nucleocytoplasmic transport, with high representation of ribonucleoprotein biogenesis and non-coding RNA processes. These findings are indicative of vascular cell stress responses, including DNA damage responses, which may have their origins in the trauma associated with surgical harvesting and could impact the process of vascular adaptation as well as neointima proliferation. Gene ontology analysis of the downregulated genes was almost exclusively associated with actin-modulation and muscle contraction processes, which reaffirm the importance of VSMC dysfunction that occurs during neointima proliferation. These findings suggest that while this model of neointima formation recapitulates changes in previously well-characterized pathways, it generates novel insight into the complex molecular and cellular mechanisms which underpin neointima proliferation.

Proteomics profiling largely corroborated both the histologic and RNA-seq findings, demonstrating dynamic changes in expressed proteins, with large alterations between the Day 0 and Day 7 SV. Actin-modulating and contractile proteins were the most highly represented in the dataset, yet untargeted proteomic analysis did not show significant changes between Day 0 and Day 7 SV. Further validation through targeted proteomics of the contractile proteins show significant downregulation, yet not as dramatic as the changes in mRNA expression. This finding implies temporal divergence between the transcriptomic and proteomic signatures during the progression of neointima proliferation, from its initial to later phases. Blood coagulation and thrombosis, which are believed to contribute to the early phase of vein graft failure^7–9^, were the top processes represented in the gene ontology analysis. ECM-producing proteins were also significantly increased at Day 7, which suggest that elements of neointima formation are shared with other contributors to vein graft failure, including thrombosis. While we did not see a strong representation of cytokines or other inflammatory signaling proteins, we detected dysregulation of proteins which regulate mitochondrial metabolism, which has been reported to occur in the presence of cellular stress responses and DNA damage^44,45^. Given the high transcriptomic and proteomic signature of VSMC, this suggests that perturbations in mitochondrial metabolism may accompany VSMC phenotypic changes during neointima proliferation.

Spatial transcriptomics is a relatively new technique that profiles gene expression at distinct locations within the blood vessel wall, thus enabling the identification of cellular subpopulations inhabiting the neointima. We first examined the difference in gene expression between Day 0 and Day 7 SV. While Day 0 SV exhibited few clusters with similar gene expression, Day 7 SV showed many more clusters which had significantly altered gene expression profiles. The cellular composition of the SV at both Day 0 and Day 7 was predicted to be a mixture of fibroblasts and VSMC, with no representation of EC or pericytes. Day 0 SV were primarily composed of healthy, contractile VSMC and secretory fibroblasts. Surprisingly, more fibroblasts than VSMC were located in the tunica media at Day 0. Additionally, the intima of the Day 0 SV was exclusively composed of transdifferentiating VSMC, which suggests that initial processes of neointima formation, including VSMC fate switching and migration, are already active at the time of SV implantation. While at Day 7, the intima was still composed of mostly fibroblasts and VSMC, UMAP showed that the gene expression profiles shared almost no overlap from Day 0.

The secretory fibroblasts predominant at Day 0, with high expression of genes related to ECM-forming, including *ELN, DCN, COL3A1*, were replaced by degradative fibroblasts at Day 7, including *MMP2, MMP14* and *TIMP1* in Day 7 SV. These degradative fibroblasts appear to remain largely in the adventitia and the outer media and may contribute to the significant fragmentation of elastin detected at Day 7. However, these fibroblasts do not appear to migrate into the neointima, where significant ECM secretion and deposition is occurring. Instead, the neointima is dominated by highly dysregulated VSMC which appear to have transdifferentitated into many fates, including previously reported synthetic, *PCNA+* VSMC. We also report novel VSMC clusters, including osteogenic VSMC, which express retinoid-induced genes including *AKAP12*, and *SPP1^+^, LGALS3^+^* VSMC, which primarily reside within the neointima. While most of the transdifferentiated VSMC are *PCNA+* synthetic VSMC, the two novel VSMC cell populations reflect the remarkable plasticity of VSMC, which can undergo various transformations in response to different pathological stimulus.

While this is the first detailed study to characterize the molecular and cellular processes involved in neointima proliferation in human blood vessels, it has several noteworthy limitations. First, given the small size of the vein pieces, we are unable to apply pressure and flow, which are important determinants of both vascular adaptation and neointima proliferation^46,47^. Second, our system is devoid of humoral and cellular components of the immune system, which is an important factor involved in SV graft disease. Third, the cohort size was insufficient to permit a complete characterization of neointima proliferation and its association with clinical variables, and we only studied two points in time (Day 0 and Day 7), which provides limited data with regard to the cellular and molecular dynamics of neointima proliferation. Fourth, weak representation of EC signatures was observed in all of the sequencing studies, including spatial transcriptomics. This likely reflects the limited resolution inherent to the current spatial transcriptomics technology, which prevents the definitive identification of these cell clusters, In conclusion, we have interrogated the process of neointima proliferation in human SV using multi-omics analyses. We detected significant sex differences in both the neointima and vessel area, as well as a negative association between BMI and neointima proliferation that was observed only in men. Our molecular and proteomic analyses point towards induction of vascular cell stress and metabolic responses that may be initiated by trauma associated with surgical procurement, as well as increases in coagulation and thrombosis that could contribute to early graft occlusion. Finally, using spatial transcriptomics, we have detected novel fibroblast and VSMC populations that that are dynamically activated and may contribute to the elastin degradation and neointimal proliferation characteristic of this model.

## Acknowledgment

This study was funded by grants NIH AG076235 (N.L.W), AHA 971459 (N.L.W) and AHA 863622 (N.L.W).

## Notes

### Competing Interest Statement

The authors have declared no competing interest.

### Summary of Updates

Mistake has been corrected by removing Joseph M. Miano from the author list.

